# A COJEC-chemotherapy resistant model of Th-*ALK(F1174L)/MYCN* neuroblastoma offers insights into tumour immune evasion and development of the bone marrow metastatic niche

**DOI:** 10.64898/2026.09.01.743331

**Authors:** Elizabeth Ruth Tucker, Andrea Lampis, Jacob Househam, Duncan Roberts, Karen Barker, Barbara Martins da Costa, Kevin Greenslade, Sarah Mayfield, Federica Lorenzi, Emma Thorley, Vidur Tandon, Nan Sophia Han, Giulia Emanuelli, Marwane Bourdim, Georgia Martin, Nicholas Willumsen, Marco Pannone, Jeremy Pryce, Paola Angelini, Mariia Yuneva, Ninib Baryawno, Trevor Graham, Sally George, Louis Chesler

## Abstract

Multi-agent COJEC chemotherapy is the main-stay of induction treatment for patients diagnosed with high-risk neuroblastoma. However, at least 10% of patients will be primary refractory to chemotherapy and only 50% achieve 5-year overall survival. The bone marrow is the most frequent site of metastasis in these patients. Novel approaches are required to improve response rates but the inter- and intra-patient tumour heterogeneity and dynamics of the neuroblastoma immune microenvironment makes anticipation of resistance phenotypes incredibly challenging.

We present here a novel immunocompetent C57 Bl/6 model of Th-*ALK(F1174L)/MYCN* neuroblastoma, in which spontaneous abdominal tumours are driven by expression of mutant Anaplastic Lymphoma Kinase and over-expression of Mycn in the neural crest. We have used this model to generate a personalised dosing schedule inducing COJEC-chemotherapy resistance, in which individual mice receive chemotherapy cycles dependent upon the progression of their neuroblastoma tumours. Using both scRNAseq and spatial immunophenotyping gave us extraordinary precision in our comprehensive analysis of the tumour intrinsic and microenvironmental factors associated with COJEC resistance. We found that the resistance phenotype was driven by *Cdk8* upregulation in adrenergic and mesenchymal tumour cells. Infiltration of immunosuppressive myeloid-derived immune cells and remodeling of the tumour-associated stroma further contributed to COJEC resistance. In the bone marrow we observed expansion of neutrophils and evidence of NETosis associated with micro-metastatic disease.

Our results further endorse the development of CDK8-targeting therapeutics for neuroblastoma patients which might boost the anti-tumour immune response. Additional studies will be required to define the roles of neutrophils and neutrophil NETosis in neuroblastoma progression and metastasis. Our C57 Bl/6 model will be pivotal in future preclinical studies of immune-modulating therapeutics.

## Background

Neuroblastoma is a paediatric malignancy of the sympathetic nervous system, commonly presenting as an abdominal mass arising from sympathetic ganglia, including the adrenal medulla. Children with high-risk neuroblastoma are frequently diagnosed with wide-spread metastatic disease involving the bone and bone marrow. Multi-agent chemotherapy is the mainstay of induction treatment, with the aim of debulking the primary tumour and diminishing metastases. In Europe, COJEC chemotherapy, consisting of cisplatin (C), vincristine (O), carboplatin (J), etoposide (E) and cyclophosphamide (C), is administered rapidly to enable a subsequent attempt at surgical resection and consolidation therapy (1). Despite intensive COJEC chemotherapy only 70-80% of patients achieve an adequate metastatic response at the end of induction, and five-year overall survival is only 50% (2, 3). 10% of high-risk neuroblastoma tumours progress through induction therapy, predicting a dismal prognosis.

Neuroblastoma tumour characteristics associated with inferior survival include amplification of the *MYCN* oncogene and Anaplastic Lymphoma Kinase (*ALK*) mutation or amplification (4, 5). Novel therapy incorporated into standard treatment takes advantage of Disialogangioside (GD2) tumour-associated antigen expression which is largely restricted to neuroblastoma cells, with anti-GD2 immunotherapy now administered during consolidation and at relapse (6). The small molecule ALK inhibitor, lorlatinib, has shown some activity in early phase clinical studies for patients with relapsing ALK mutant neuroblastoma, and it is now being evaluated as an addition to induction therapy (7). However, neuroblastoma tumour heterogeneity and phenotypic dynamics are limiting the translational benefit and durable response of novel therapies such as these (8, 9). Neuroblastoma cell lineage state, which exhibits plasticity in some model systems, can result in downregulation of GD2 (6) and ALK (10) in mesenchymal-type cells. Transition from adrenergic lineage to mesenchymal is also associated with neuroblastoma chemotherapy resistance and relapse, which has recently been shown to promote immunogenicity through the epigenetic regulation of inflammatory and immune response genes (11).

The immune-suppressive and -evasive microenvironment of neuroblastoma is the result of low mutational burden and resultant scarcity of neoepitopes (12). *MYCN* amplification, found in 50% of high-risk neuroblastoma tumours, leads to downregulation of major histocompatibility complex (MHC) class one molecules(13).

Additionally, the microenvironment is infiltrated by immunosuppressive tumour-associated macrophages (TAMs) and cancer-associated fibroblasts (CAFs), with exclusion of T cells, B cells and NK cells. There is a huge drive to develop immunotherapy options for neuroblastoma patients, through the discovery of neuroblastoma-unique immune vulnerabilities. This includes targeting the NECTIN-TIGIT and PD-L1 checkpoints which together achieved synergy in a syngeneic allograft of the chemotherapy-resistant “TAM6” Th-*ALK(F1174L)/MYCN* 129svj model (14).

However, there remains a paucity of preclinical immunocompetent models with which to study multi-chemotherapy-resistant neuroblastoma. Moreover, the existing genetically engineered murine models (GEMM) of neuroblastoma do not develop overt metastatic disease. Transgenic models remain an essential component of the preclinical toolkit for dissecting the biology of these rare tumours and evaluating novel therapeutic approaches in tumour immune microenvironment. The Th-*MYCN* GEMM develops abdominal neuroblastoma tumours spontaneously through over expression of *MYCN* in neuroectodermal cells, with gains and losses of chromosomes in regions syntenic with human neuroblastoma(15). scRNAseq has revealed that Th-*MYCN* tumours consist of a variety of cell types, with tumour cells arising from both adrenergic sympathoblasts and chromaffin cells and share transcriptional profiles with human neuroblastomas (16). Double heterozygous Th-*ALK(F1174L)/MYCN* GEMM tumours have demonstrated responses to small molecule inhibitors of ALK, reflecting what has been observed in early phase clinical trials (17, 18). Both the Th-*MYCN* and *Th-ALK(F1174L)/MYCN* GEMMs were originally derived in the 129svj background strain, in which they are highly penetrant, but tumour immunological research is optimally conducted in C57 Bl/6 mice, which have enhanced cell-mediated immunity (19). A recently described *Mycn*-driven, murine neural crest-derived, transplantable nGEMM model of neuroblastoma was generated in the Bl/6 strain (20). This model represented immune-cold human neuroblastoma with a predominance of myeloid cells and immunosuppressive TAMs.

We induced resistance to induction COJEC chemotherapy in a fully back-crossed 10^th^ generation C57 Bl/6 Th-*ALK(F1174L)/MYCN* model of high-risk neuroblastoma, to elucidate clinically-relevant tumour-intrinsic and immune microenvironmental features associated with multi-agent chemotherapy resistance. Our unique model emphasises the role of myeloid-derived immunosuppression in chemotherapy resistance, driven by upregulation of *cdk8*. Furthermore we observe stromal remodelling as a feature of the immunosuppressive resistant tumour microenvironment, with expansion of neutrophils and NETosis associated with micro-metastatic spread to the bone marrow.

## Results

### Generation of COJEC chemotherapy-resistant C57 Bl/6 Th-*ALK(F1174L)/MYCN* mice

The Th-*ALK(F1174L)/MYCN* GEMM was originally generated in the 129svj tumour-permissive strain (21). We bred 129svj Th-*MYCN* and Th-*ALK(F1174L)* mice sequentially with wild-type C57 Bl/6 mice, until we reached the 10^th^ generation backcross. At that point we bred single heterozygous Th-*MYCN* and Th-*ALK(F1174L)* to generate double heterozygous C57 Bl/6 Th-*ALK(F1174L)/MYCN*. We saw a significantly increased latency in C57 Bl/6 and double heterozygous mice had 100% abdominal tumour penetrance by 150 days (figure 1a). Th-*ALK(F1174L)* heterozygous mice had no phenotype, as also seen in the 129svj model. Heterozygous and homozygous Th-*MYCN* mice had a zero and <10% tumour penetrance respectively (supplementary figure 1a). Retrospective analysis of mendelian ratios suggested that homozygous Th-*MYCN* in the C57 Bl/6 background were less viable *in utero* (data not shown).

**Figure One:**
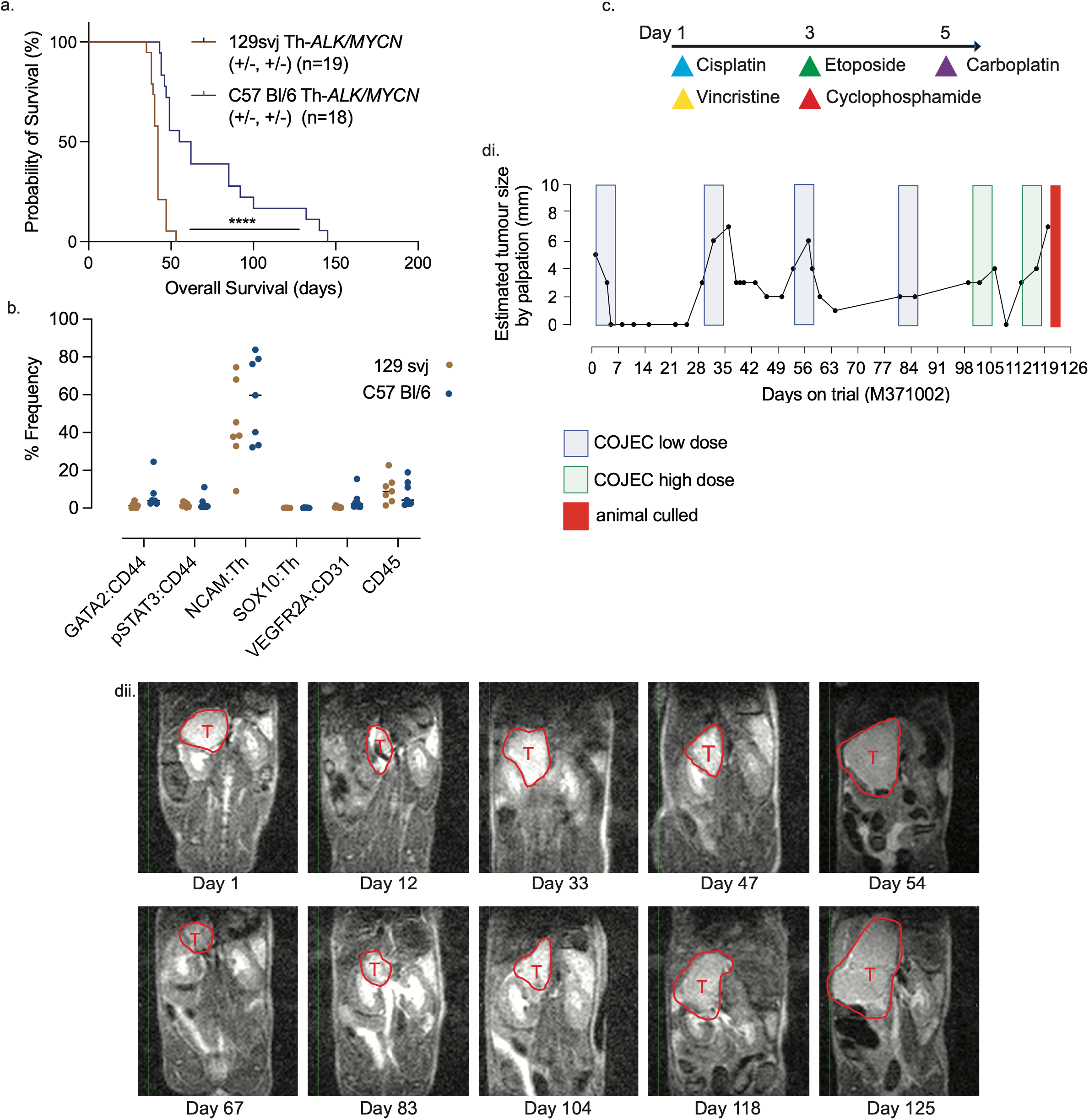
Generation of COJEC chemotherapy-resistant C57 Bl/6 Th- *ALK(F1174L)/MYCN* neuroblastoma mice. a. Survival of C57 Bl/6 versus 129 svj Th-*ALK(F1174L)/MYCN* mice. b. Immunofluoresecence (Opal) quantification demonstrates no significant differences between C57 Bl/6 and 129svj Th-*ALK(F1174L)/MYCN* tumours at ethical endpoint. c. Schematic of COJEC chemotherapy treatment cycle. d. Tumour palpation data from one Th-*ALK(F1174L)/MYCN* mouse treated with cycles of COJEC chemotherapy according to a personalised dosing schedule. e. MRI of mouse depicted in d to show a single primary tumour (T, outlined in red) progressing and regressing during cycles of COJEC.

Histopathological comparison of Th-*ALK(F1174L)/MYCN* tumours in the 129 svj and Bl/6 strains did not reveal any significantly different expression of markers associated with either adrenergic or mesenchymal status, or with immune cell or vascular phenotype at ethical endpoint (figure 1b and supplementary figure 1b). However, the C57 Bl/6 mice exhibited a superior response to the ALK inhibitor lorlatinib, when treatment was initiated at an intermediate abdominal tumour size of approximately 5mm (supplementary figure 1ci-ii). Altogether, our initial strain comparison suggested that Th-*ALK(F1174L)/MYCN* tumours arising in the Bl/6 background retained the neuroblastoma phenotype similarily to 129 svj background tumours. However, the enhanced cell-mediated immunity of the C57 Bl/6 was likely to be supporting the improved efficacy to ALK inhibition. We next used C57 Bl/6 Th-*ALK(F1174L)/MYCN* to model neuroblastoma induction chemotherapy resistance. We adopted the murine dosing schedule for COJEC chemotherapy, originally described by Mañas *et al* (22) , but with doses lowered to account for the lesser drug tolerability of immunocompetent animals. Our treatment strategy to induce resistance consisted of cycles of COJEC administered over five days, and following the first cycle, animals were only treated with subsequent cycles of chemotherapy when tumour progression re-occured (figure 1c). In total, we treated 12 animals with cycles of COJEC to become resistant and each animal required a different number and timing of cycles (figure 1d and supplementary figure 2). Magnetic Resonance Imaging (MRI) was also used to monitor tumour growth and regression more accurately when palpation size was indeterminant (figure 1e).

### Single cell RNA sequencing of tumours suggests *Cdk8* is mediating both the tumour-intrinsic and immune inflammatory COJEC-resistant phenotype

To elucidate the interplay between tumour -intrinsic and -extrinsic mechanisms leading to resistance, we undertook scRNAseq of 7 COJEC-resistant Th-*ALK(F1174L)/MYCN* tumours and 8 vehicle-treated tumours, alongside bone marrow from the femurs of each animal. To assist in the accurate annotation of tumour cells and to investigate the presence of bone marrow neuroblastoma cell metastases, we additionally undertook scRNAseq of 4 bone marrows taken from the femurs of wild-type (WT) C57 Bl/6 animals, aged matched for COJEC-resistant mice.

Cell type annotation enabled us to clearly identify clusters previously characterised by RNAseq of both human and murine neuroblastomas (figure 2a and supplementary figure 3a). We identified both adrenergic and mesenchymal neuroblastoma tumour cells in both vehicle control and COJEC-resistant tumours, which did not significantly shift in frequency between treatment groups (supplementary figure 3b), and separately, neural progenitors. The proportion of Schwann Stroma Cells was not significantly enriched in COJEC-resistant tumours (supplementary figure 3c), consistent with recent data comparing Schwannian stroma abundance in high-risk clinical neuroblastoma tumours at the time of surgical resection, where there was no significant increase in stroma from *MYCN*-amplified tumours (23).

**Figure Two:**
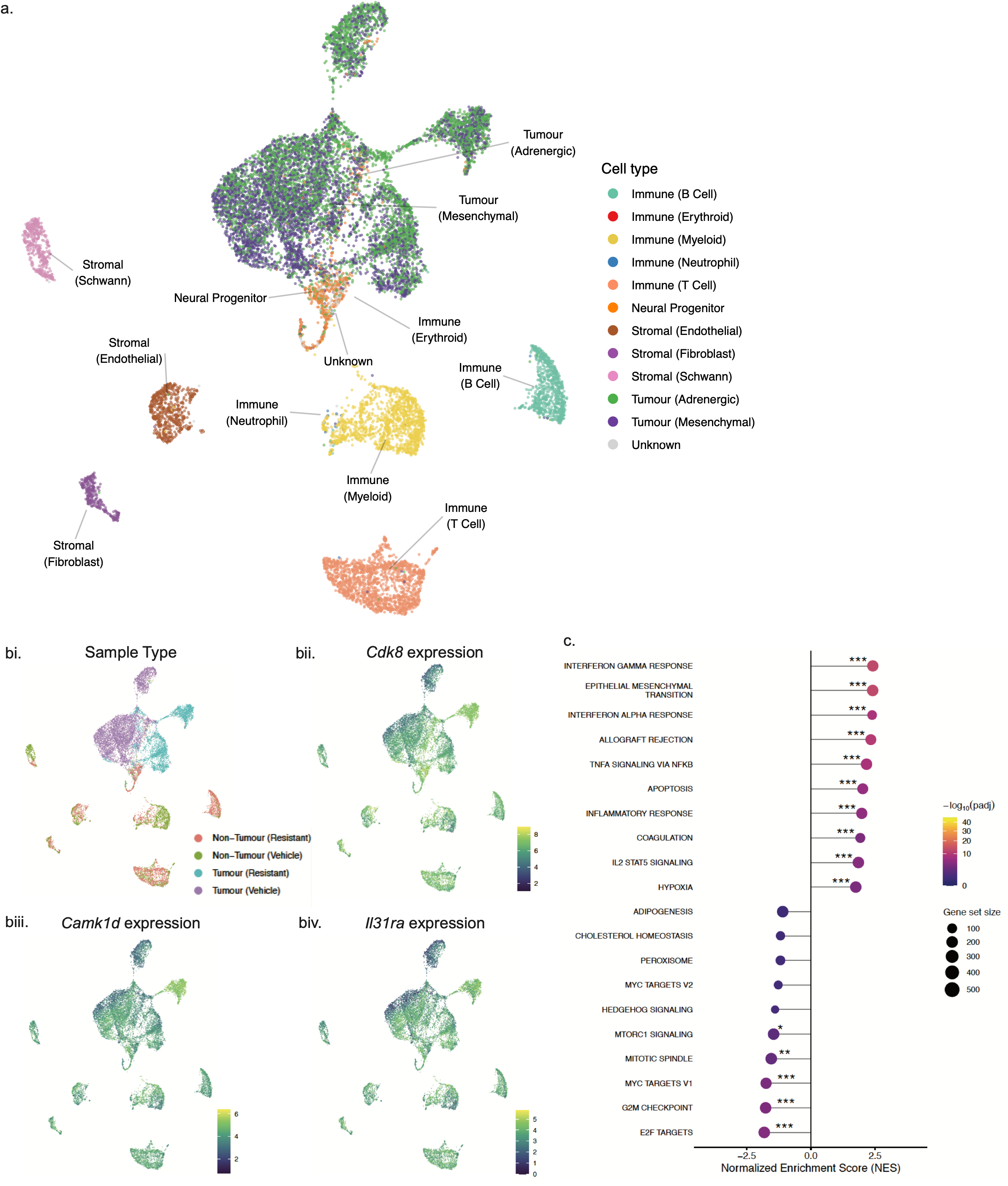
Single cell RNA sequencing of tumours suggests Cdk8 is mediating both the tumour-intrinsic and immune inflammatory COJEC-resistant phenotype. a. UMAP of scRNAseq with all samples overlaid (COJEC-resistant Th-*ALK(F1174L)/MYCN* tumours n=7, vehicle control tumours n=8). bi. UMAP of sample group from tumour samples. UMAP illustrating gene expression of *Cdk8* (bii), *Camk1d* (biii) and *Il31ra* (biv). c. Hallmark GSEA comparing COJEC-resistant and vehicle control tumours.

The comparison of sample groups (COJEC resistant versus vehicle) revealed distinctive populations of tumour cells restricted to either vehicle-treated or COJEC-resistant samples (figure 2bi), and deeper interrogation of the differential genes driving these phenotypes showed expression of Cdk8, activity of which is associated with reprogramming of the microenvironment in chemo-resistant cancer (24). Alongside Cdk8, we found upregulation of Camk1d (a modulator of immune resistance associated with a poor response to PD-L1 checkpoint inhibition (25)) and IL31-ra (an inflammatory intermediatory, typically associated with a type 2 allergic immune response (26)) highly upregulated in resistant samples (figure 2bii-iv, and supplementary figure 3d).

Hallmark Gene Set Enrichment Analysis (GSEA) revealed high activity of Interferon-_Y_ (IFN-_Y_), Interferon-lZ (IFN-lZ) and IL2 STAT5 signaling, indicating that inflammation was driving the immune response; attracting T cells into the tumour microenvironment (figure 2c). Alternatively, chronic exposure of the tumour microenvironment to IFN-_Y_ could be inducing immunosuppressive cells. Upregulated “TNFlZ Signaling via NF κβ” alongside downregulated “G2M checkpoint” (and an increase in the proportion of G1 tumour cells (supplementary figure 3e)) in resistant tumours, raised the possibility that they had acquired the well-described Senescence Associated Secretory Phenotype (SASP), driving an immunosuppressive myeloid-response. Pro-inflammatory SASP is well-known to result in complex cross-talk between NF- κβ, p38 MAPK, mTOR and cyclic-GMP-AMP synthase-stimulator of interferon genes (cGAS-STING) in patient-derived tumour data sets, and in alternative Bl/6 preclinical models of adult cancers (27, 28). A recent paired scRNAseq analysis of neuroblastoma patient tumour tissue from diagnosis and post-chemotherapy, also showed subsets of persister cells which had entered a non-cycling state, characterised by suppression of MYC(N) activity and concomitant high NF-κβ (23). In agreement with this, our preclinical COJEC resistant tumours showed downregulation of “MYC target v1”, “MYC target v2” and “MTORC signaling”.

### Immunosuppressive myeloid populations drive COJEC resistance

To determine the origins of cytokines and chemokines which might be driving the immunosuppressive microenvironment in our resistant model, we interrogated and combined two preclinical SASP gene sets against our data (figure 3a) (27, 28). This analysis revealed that myeloid cells, especially neutrophils, were the most active subtypes of immune cells within resistant tumours. Several markers of immunosuppressive tumour-associated neutrophils (TAN) were identified, including the chemoattractants *Csf1*, *Cxcl2* and *Tgfb1*, and other SASP markers, including *Cdkn1a*, *Cdk15*, *Olr1*, *Ramp1*, *Ptgs2*, *Itgax*, *Lmnb1* and *Fes*. *Ramp1* is especially associated with facilitation of the neuro-immune connection to myeloid cells (29). *Cxcr2* was upregulated in myeloid cells (not including neutrophils), strongly suggesting that the Cxcl2-Cxcr2 axis was involved in the recruitment of myeloid-derived suppressor cells (MDSC) into the tumours. Notably, in primary senescent stroma, there was upregulation of two members of the *Serpine* gene family (*Serpine1* in Schwann stroma and *Sepina9* in fibroblast stroma) which have been associated with remodelling of the extracellular matrix in a model of gastric cancer, an inflammatory mediator in pancreatic ductal adenocarcinoma, and in neuroblastoma, have been associated with high-risk poor prognosis disease in metabolomics data (30–32).

**Figure Three:**
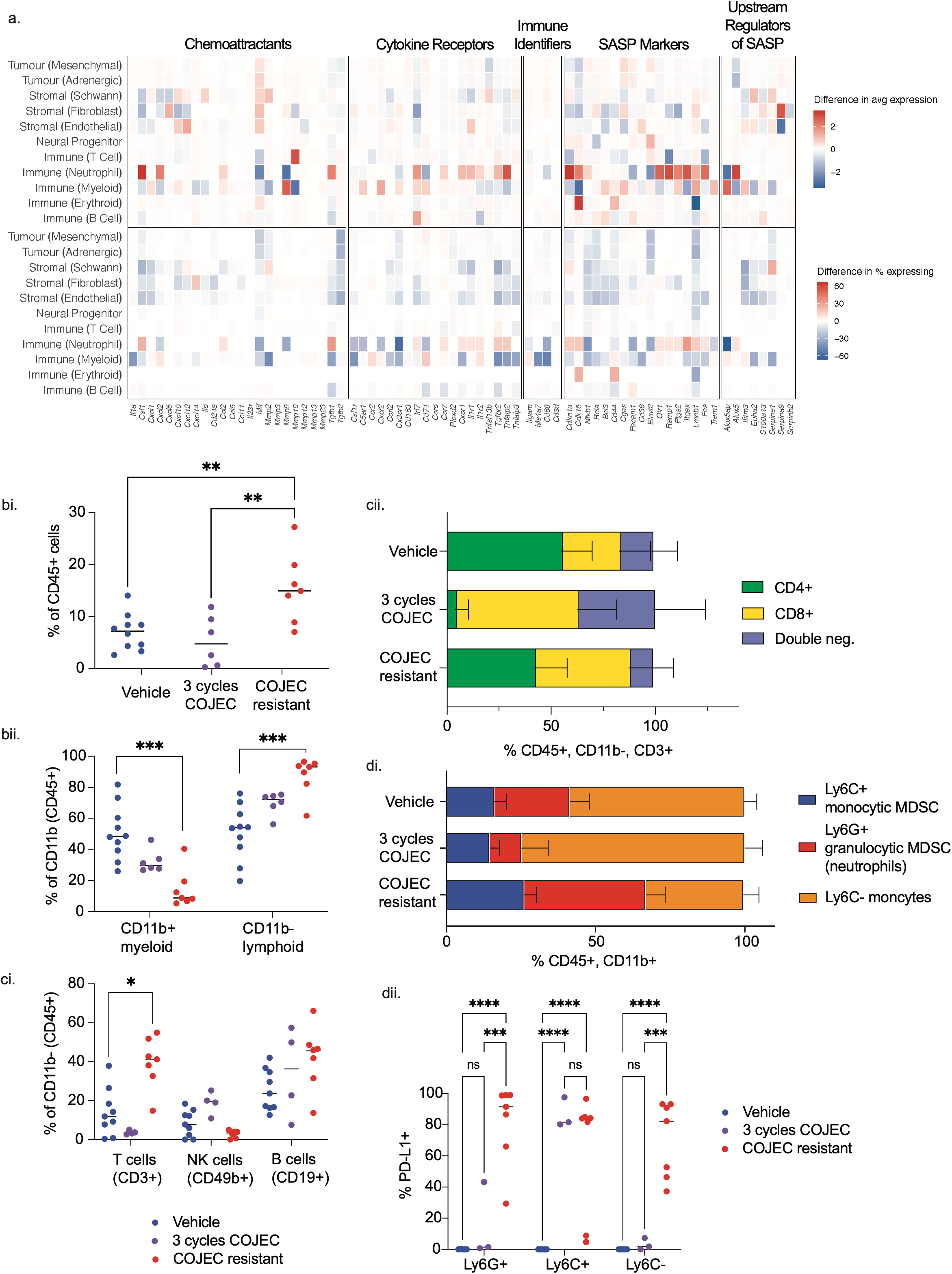
Tumour-associated neutrophils are driving an immunosuppressive tumour microenvironment in COJEC chemotherapy-resistant C57 Bl/6 Th- *ALK(F1174L)/MYCN*. a. Bubble plot of combined SASP signatures showing gene marker expression. Dot size indicates fraction of expressing cells, colour indicates Z-score-normalized expression (27, 28). b-d. Flow cytometry panel analysis of tumours from vehicle control mice, mice treated with 3 cycles of COJEC chemotherapy, and COJEC-resistant mice. b.i. % of CD45+ cells. b.ii. % of CD11b+ and CD11b-(CD45+) cells. c.i. CD11b- lymphocyte subsets (CD3+, CD49b+ and CD19+). c.ii. Proportion of CD4+, CD8+ and double negative CD3+ T cells. d.i. Proportion of Ly6C+, Ly6G+ and Ly6C-, CD11b+ myeloid cells. d.ii. PD-L1 positivity of myeloid subsets.

Flow cytometric analysis of tumours included a third cohort of tumours from mice which had been treated with only 3 cycles of COJEC before samples were taken, to monitor shifts in immune populations prior to the development of resistance. There was an overall increase in CD45+ immune cells infiltrating COJEC-resistant tumours (figure 3bi), accounted for by an increase in CD11b- lymphoid cells, especially CD3+, and a relative reduction in CD11b+ myeloid cells (figures 3b and c). There were no markers of T-cell activity or exhaustion at the COJEC-resistant endpoint, but after three cycles of COJEC, Ki67 and PD-1 positivity was noted on CD8+ cytotoxic T-cells, suggesting that COJEC-resistance was associated with a loss of CD8+ activity in Th-*ALK(F1174L)/MYCN* tumours (supplementary figures 4a and b). Although there was a reduction in the number of CD11b+ MDSC and monocytes in COJEC resistant tumours, there was re-expansion of Ly6G+ PMN-MDSC (polymorphonuclear myeloid-derived suppressor cells) neutrophils at the COJEC resistant experimental endpoint (figure 3di). All myeloid populations were noted to be strongly enriched for PD-L1 positivity (figure 3dii), consistent with the scRNAseq data in demonstrating intense immunosuppressive activity of MDSC in COJEC-resistant tumours. Notably, we observed a low and inconsistent positivity for PD-L1 and NECTIN on tumour cells (identified by GD2 positive staining), which is in line with emerging clinical observations that tumours with predominant myeloid immunosuppression characterise a sub-group of non-responders to check-point inhibition therapy (data not shown) (33).

Altogether our results show activation of MDSC, and especially PMN-MDSC/neutrophils, in COJEC-resistant Th-*ALK(F1174L)/MYCN* tumours, despite the relative decrease in myeloid cell numbers in the tumour microenvironment.

### Stromal remodelling and immune infiltration of COJEC-resistant tumours

We next used spatial immunophenotyping to characterise the location of tumour and immune cell population subtypes associated with the acquisition of COJEC resistance. Following annotation, immune cell counts were consistent with the preceding flow cytometry analysis, with a significant increase in PMN-MDSC neutrophil (Ly6G+) and additionally, increased M2 macrophage (CD206+) frequency in COJEC-resistant tumours (supplementary figures 5ai (lymphoid) and ii (myeloid)). Also observed was a significant decrease in dendritic cell (CD11c+) frequency in COJEC-resistant tumours, suggesting immune suppression via a reduction in antigen presentation to T-cells also contributed to tumour progression, and was consistent with reduced Ki67+ and PD-1+ CD8+ T-cells in flow cytometry data. Double negative T-cells were increased in frequency on the spatial analysis between vehicle and COJEC-resistant tumours, raising the possility that _Y_δ T-cell infiltration was an additional mechanism of immune suppression in this model. The density of CD44+ stroma (not discriminating between schwannian and fibroblastic stroma) was not significantly altered between vehicle and COJEC-resistant tumours (supplementary figure 5b).

Neighbourhood enrichment analysis demonstrated the remodelling of tumour immune populations and stromal architecture during the evolution towards COJEC-resistance (figures 4ai (vehicle tumours) and 4aii (COJEC-resistant tumours), with bi and bii, example neighbourhood ID maps of COJEC-resistant tumours) (34). Both vehicle and resistant tumours showed immune hot regions (cluster 4 and cluster 3 respectively) and immune-depleted tumour regions (clusters 3 and 1 respectively). A stromal neighbourhood with immune infiltration enriched with M1 and M2 macrophages, monocytes and neutrophils, was unique to the COJEC-resistant tumours (Cluster 0 figure 4aii, figure 4ci (phenocycler). This was further evidenced spatially by topographical correlation analysis showing significant positive correlation of stroma and M2 macrophages, thus suggesting high colocalization within 80µm (figure cii (dot plot), ciii and civ (topographical correlation maps), supplementary figure 5c). In contrast, stroma in the vehicle-treated tumours was largely immune depleted, except for M1 macrophages. Spatial colocalization of M2 macrophages within stromal enriched boundary regions supported the scRNAseq and flow cytometry data suggesting MDSC-mediated immunosuppression in COJEC-resistant Th-*ALK(F1174L)/MYCN* tumours, with evidence of a physical barrier being created by stromal cells alluding to additional immune exclusion tactics by COJEC-resistant tumours.

**Figure Four:**
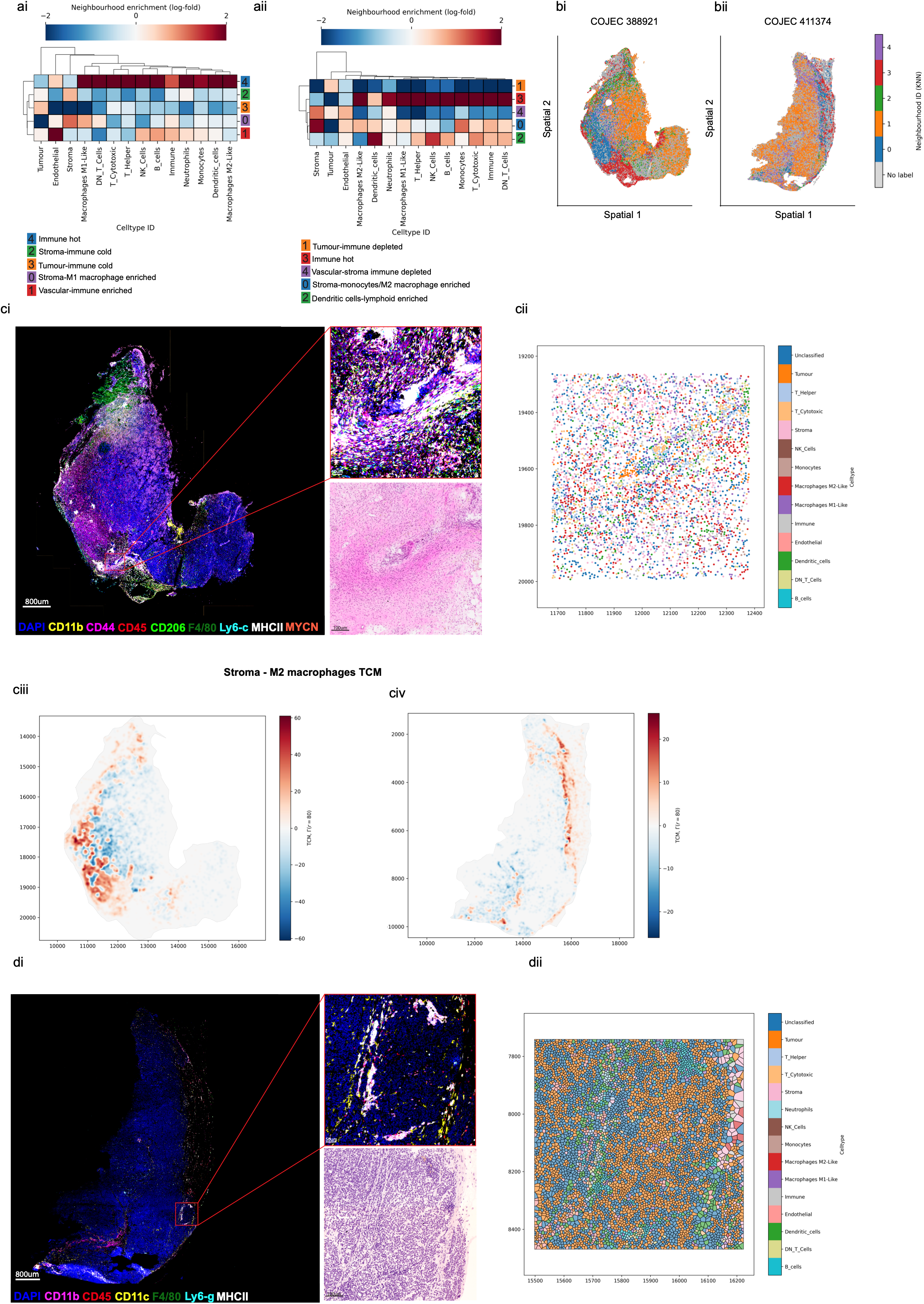
Stromal remodelling and immune infiltration is associated with the acquisition of COJEC chemotherapy-resistance. Neighbourhood enrichment matrix of a.i. vehicle control tumours (n=8) and a.ii. COJEC-resistant tumours (n=8). b.i. and ii. Example neighbourhood ID maps of COJEC-resistant tumours. c.i. Example tissue sections depicting Neighbourhood Cluster 0, from COJEC-resistant tumour (a.ii) with extensive infiltration of M2 macrophages (CD45, MHCII, Ly6-c and CD206 positive) within stroma (CD44). Region of interest highlighted in red box with corresponding H&E below. c.ii. Corresponding dot-plot to highlight broad myeloid cell infiltration. c.iii. and c.iv. topographical correlation maps of COJEC-resistant tumour, emphasising leading edge of Neighbourhood 0 (Stroma Monocyte/M2 macrophage enriched) alongside tumour tissue. d.i. Example tissue section depicting Neighbourhood Cluster 2, from a COJEC-resistant tumour (a.ii). Region of interest highlighted in red box with corresponding H&E below. d.ii. Voronoi plot illustrating T-cell cluster surrounding dendritic cell hot-spot.

Despite the overall reduction in frequency of dendritic cells (CD11c) in COJEC-resistant tumours, these were noted to be enriched in regions of the tumour and associated with lymphoid immune cells (neighbourhood cluster 2, Figure 4di (phenocycler) and 4dii (Voronoi plot). Supplementary figures 5 di and dii (COJEC-resistant tumours) CD11c positive niches). This suggested that anti-tumour immunity mediated by residual dendritic cells was still active in COJEC-resistant tumours, but may have passed an equilibrium stage, whereby myeloid immunosuppressive mechanisms were favouring tumour progression in alignment with the objective tumour measurement.

A further neighbourhood cluster (Cluster 4, supplementary figures 5ei and eii) identified uniquely in COJEC-resistant tumours, consisted of stroma (CD44), endothelial cells (CD31) and neutrophils (Ly6G), raising the possibility that a distinctive sub-group of neutrophils might be supporting intravasation of tumour cells in this model.

### Establishment of the bone marrow metastatic niche

Patients with high-risk neuroblastoma most frequently present with metastases to the bone marrow, and this site is also the most common site of metastatic relapse. Therefore, we decided to investigate the frequency of spontaneous metastases to the bone marrow in our model.

Flow cytometry panel analysis of bone marrows from the femurs of vehicle treated and COJEC-resistant mice demonstrated a low frequency of neuroblastoma cells (supplementary figure 6a). There was no significant enrichment of GD2+ cells associated with COJEC-resistance (GD2+ (CD45-, CD11b-) cells in the bone marrow from vehicle treated animals ranged 0.24-3.66%, and in COJEC-resistant animals ranged 0.03-5.69%). Inclusion of bone marrow samples from four WT C57 Bl/6 mice in the scRNAseq, allowed confidence in the annotation of neuroblastoma cells in these samples, particularly as the bone marrow is a highly innervated tissue type, rich with neural progenitors (figure 5ai-ii). Adrenergic-type neuroblastoma cells were predominantly noted in the bone marrow of Th-*ALK(F1174L)/MYCN* animals, and these were enriched in COJEC-resistant mice. However, the overall number of cells sequenced was 10x lower than that for the tumour samples, which may have accounted for the lack of any significant findings in the differential gene expression (DEG) analysis between vehicle and resistant bone marrow samples (supplementary figure 6b). However, DEG comparison between wild-type bone marrow and bone marrow from vehicle-treated mice revealed the upregulation of several pro-metastatic genes, including *Aff3*, *Hnrnpul1*, *Ctsd*, *Kpna1* and *Arhgef1* (supplementary figure 6 ci-ii). ARHEF1 acts to suppress T-cell activity in metastatic sites which can be reversed by COX-1 inhibition (including aspirin treatment) to provoke immune-mediated rejection of disseminated cancer cells (35).

Despite the lack of significant changes between vehicle and COJEC-resistant scRNAseq bone samples, flow cytometry analysis demonstrated that CD11b+ myeloid cells dominated the bone marrow niche (figure 5 bi). Mice culled following 3 cycles of COJEC had a lesser proportion of CD11b+ myeloid cells in the bone marrow, consistent with chemotherapy-induced cytopenia. After 3 cycles of COJEC the Ly6G+ PMN-MDSC neutrophils were reduced, but showed a trend towards re-expansion in the COJEC-resistant mice (figure 5 bii). This population was not PD-L1 positive, in contrast to the findings in the analysis of the primary tumours, suggesting immunosuppression was not a major role of neutrophils in the bone marrow (supplementary figure 6d).

**Figure Five:**
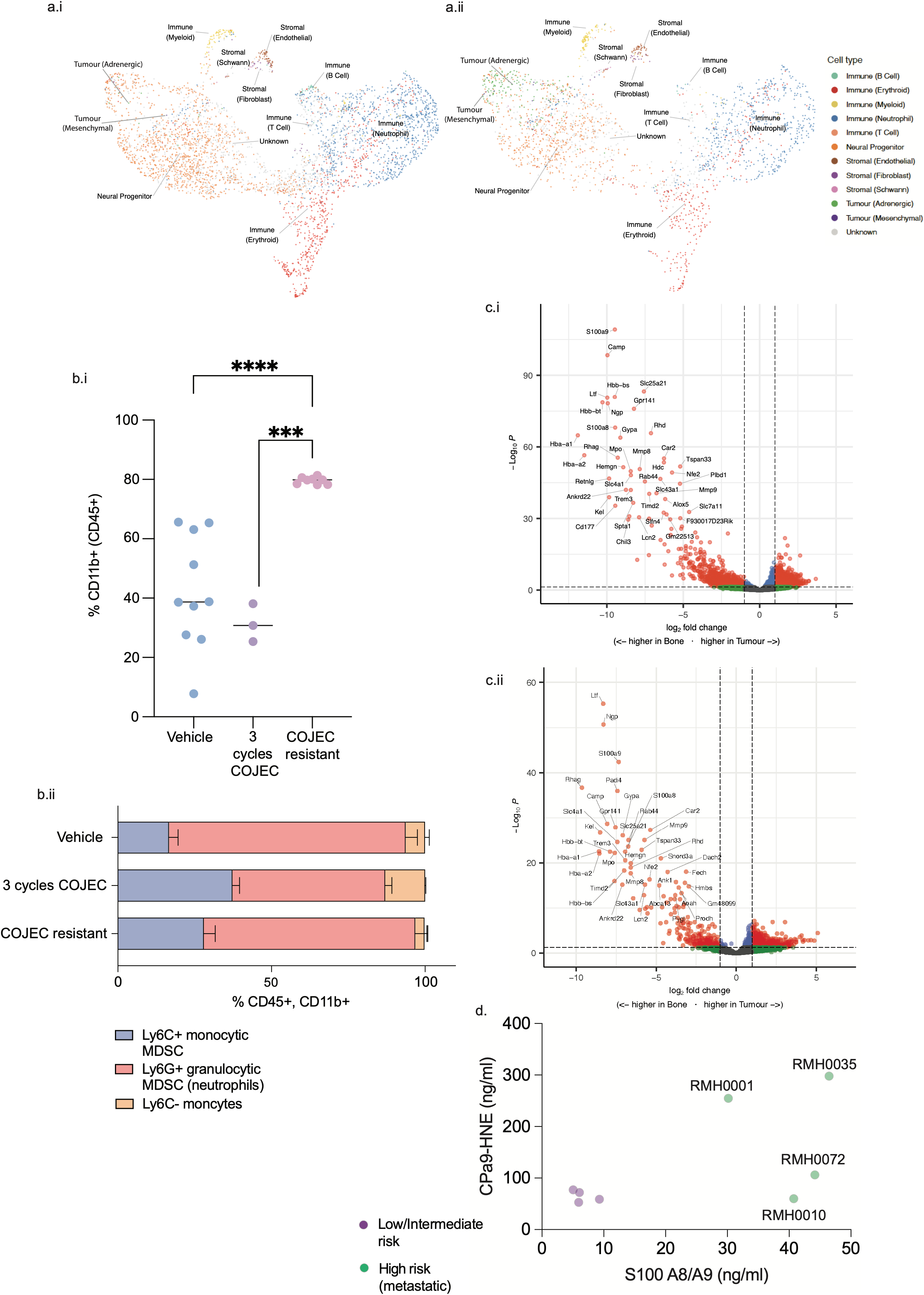
Micro metastatic disease in the bone marrow of C57 Bl/6 Th- *ALK(F1174L)/MYCN* mice is associated with re-expansion of a neutrophil population. a. Bone marrow scRNAseq UMAP from i. vehicle control mice (n=8) and ii. COJEC-resistant mice (n=7). b. Flow cytometric analysis of i. CD11b+ cells in the bone marrow and ii. CD11b+ MDSC composition in the bone marrow. c. scRNAseq DEG of paired tumour and bone from i. Vehicle control mice and ii. COJEC resistant mice. d. S100 A8/A9 and CPa9-HNE immunoassays of neuroblastoma patient plasma samples, taken at diagnosis (pre-treatment). (Purple: low/intermediate risk patients; Green: high-risk, metastatic patients)

To interrogate this further, we performed a paired comparison between the tumours and bone marrow samples via scRNAseq. The DEG highlighted several genes related to different stages of neutrophil development, and also neutrophil extracellular trap (NET) activity highly upregulated in the bone marrow *(Ltf, Ngp, Mpo, s100a8, s100a9, Mmp8, Mmp9, Padi4*) (figure 5ci-ii). Immunohistochemical staining of Neutrophil elastase and Citrullinated H3 showed strong positivity in bone marrow samples from all tumour-bearing animals (supplementary figure 6e).

Although neutrophil NETs have been reported in human neuroblastoma, their role is not well understood, due to the difficulty in their isolation and identification. The S100A8/A9 heterodimer (calprotectin) is widely used marker of neutrophil NETs in clinical inflammatory diseases, but S100 proteins are also expressed by neuroblastoma schwannian stroma. Therefore, we sought to confirm our preclinical findings, by carrying out an immunoassay specific for neutrophil-elastase cleaved S100A8/A9 (CPa9-NHE), across a panel of neuroblastoma patient diagnostic blood plasmas (figure 5d). This cohort included samples from both patients with metastatic high-risk neuroblastoma, and low-, or intermediate-risk non-metastatic neuroblastoma. Whilst all four patients in the high-risk group had elevated S100A8/A9 as expected, only two patients in this group had clinically significantly raised levels of the cleaved CPa9-HNE heterodimer at diagnosis (RMH0001 and RMH0035). These two patients were distinguished from rest of the group by their treatment-refractory extensive bone metastases. Patient RMH0001 presented with a mass arising from the mandible and was subsequently found to have a primary large right suprarenal neuroblastoma and further skeletal metastases (supplementary figure 7a). This patient was also primary refractory to induction COJEC chemotherapy, with *MYCN* amplified disease. Patient RMH0035 presented with widely disseminated Meta-iodobenzylguanidine (mIBG)-avid skeletal deposits in addition to a large neuroblastoma tumour in the right hemithorax (supplementary figure 7b). This patient had an ATRX-mutant tumour and had a poor metastatic response to induction COJEC chemotherapy. Reflecting this, the SIOPEN score for patient RMH0035 remained at 36 on mIBG imaging at diagnosis and post-induction (36). In contrast, patient RMH0010 had COJEC-refractory high-risk neuroblastoma with liver metastases. The skeletal metastases in this patient were associated with a SIOPEN score (calculated from mIBG scans) of 36 at induction, which reduced to 24 post-induction, and further reduced to 17 during the course of palliative treatment. Patient RMH0072, with *MYCN*-amplified disease, had an adequate response to COJEC and completed the high-risk treatment protocol. This patient had a SIOPEN score (calculated from mIBG scans) of 1 at diagnosis and post-induction, which reduced to 0 by four months into treatment.

## Discussion

COJEC chemotherapy induction was developed to administer rapidly as a life-saving intervention to children with high-risk neuroblastoma, reducing tumour burden and enabling subsequent resection of the residual primary tumour. However, primary-refractory disease and chemotherapy-resistant relapse remain significant clinical challenges. Preclinical *in vivo* modelling of COJEC chemotherapeutics can contribute an insight into tumour-intrinsic and microenvironmental resistance mechanism associated with COJEC resistance.

We have developed a COJEC-chemotherapy-resistant preclinical model of high-risk neuroblastoma, using the Th-*ALK(F1174L)/MYCN* GEMM in a C57 Bl/6 background strain. Despite the lower threshold for T cell activation and NK cell activity in Bl/6 mice, we have found that the myeloid immune compartment is driving immunosuppression and enabling neuroblastoma tumour progression in this model. However, evidence of dendritic cell neighbourhood clusters in COJEC-resistant tumours could imply residual anti-tumour immunity via antigen expression and attraction of cytotoxic T-cells, which might be responsive to checkpoint re-activation.

Although we did not observe enrichment of mesenchymal-type neuroblastoma cells in our COJEC-resistant tumours, we found distinct upregulation of *Cdk8* in this cohort across both adrenergic and mesenchymal tumour cell compartments. Cell cycle proteins are well-known to be dysregulated in malignant cells and CDK8 activity has specifically been linked to the upregulation of the IFN-_Y_ response and cytokine activity via STAT1 (37). This is in keeping with the findings in our model, in which IFN-_Y_ response was the top up-regulated gene-set in COJEC-resistant tumours.

In neuroblastoma, targeting of CDK8 sensitises *RAS*-mutant neuroblastoma cells to MEK inhibitors *in vitro* (38). Therefore, the development of CDK8 inhibitors for neuroblastoma, in which *RAS* pathway mutations are a frequent mechanism of therapeutic resistance, already has a clear rationale. *RAS*-pathway mutations are associated with ALK inhibitor resistance (39, 40), and therefore, as an ALK-driven neuroblastoma model, Th-*ALK(F1174L)/MYCN* would be an ideal *in vivo* platform to conduct further studies of CDK8 inhibitor combinations.

CDK8 dysregulation has furthermore been associated with transcriptional activity of serum-response genes, via RNA polymerase II elongation (41). Whilst the Serum Response Factor (SRF) is ubiquitous across many cell types, notably, myeloid cells are well described to be dependent upon the SRF for their inflammatory response, including cytoskeletal conformational changes via integrin redistribution (42). Manipulation of CDK8 expression would be required to confirm whether this mechanism was primarily responsible for myeloid-derived immunosuppression in our COJEC-resistant model, but nevertheless published evidence strongly supports the association.

Intriguingly, our data also demonstrates the co-expression of *Il31ra* alongside *Cdk8* in COJEC-resistant tumour cells. IL31RA has recently been described as an intermediatory between sensory neurons and type 2 inflammation, associated with allergic disease (26). In this study, activation of IL31RA led to release of CGRP. Similarily in our model *Ramp1*, also noted to be critical for the neuro-immune connection, is upregulated in neutrophils in COJEC-resistant tumours (29). We can therefore postulate that the neuro-immune axis, a well-established physiological relationship between sympathetic neurons and the immune system, is contributing to immunosuppression in our COJEC resistant model of neuroblastoma.

We have also observed spontaneous micro-metastatic spread of neuroblastoma cells to the bone marrow in both our vehicle control and COJEC-resistant mice. The bone marrow is the most common site of metastasis in human neuroblastoma, and lack of spontaneous metastases has been cited as a weakness of transgenic models, with the exception of Th-*MYCN* mice with caspase-8 deficiency (43). In our C57 Bl/6 neuroblastoma model metastasis was associated with expansion of neutrophils across all maturation states, including evidence of neutrophil NETosis. Validation of this observation in neuroblastoma patient plasma samples has revealed that neutrophil NETosis might be associated with disseminated bone metastases, but further studies are required to establish the precise role of neutrophil NETosis in osteoclastogenesis.

## Conclusions

Altogether, our model has highlighted CDK8 as a central mediator of resistance to COJEC chemotherapy in C57 Bl/6 Th-*ALK(F1174L)/MYCN* neuroblastoma. The role of CDK8 can be linked to neuro-immune axis dysregulation resulting in myeloid-derived immunosuppression. Stromal remodelling with infiltration of M2-type macrophages was apparent in our spatial analysis, suggesting concomitant immunosuppression and immune exclusion tactics are displayed by COJEC-resistant tumours. Our model also suggests an central role of neutrophils in neuroblastoma progression, chemo-resistance and metastasis, which are thus far an under-studied immune cell type in neuroblastoma.

The myeloid-dominant immunosuppression observed in this COJEC-resistant C57 Bl/6 Th-*ALK(F1174L)/MYCN*, might represent a sub-group of neuroblastoma with resistance to checkpoint inhibitors. Potential biomarkers *camk1d* and *Il31ra*, upregulated alongside *Cdk8*, deserve further interrogation in patient datasets including both checkpoint inhibitor -responsive and -refractory neuroblastomas.

### Limitations of the study

We have investigated the molecular resistance mechanisms to chemotherapy in an immunocompetent C57 Bl/6 model of neuroblastoma, using a individualised approach to timings of chemotherapy administration, and applying doses of COJEC according to tolerability studies. Whilst our findings reflect several key features of chemotherapy resistance mechanisms observed in patients, it will be essential to validate the relevance of our novel findings using patient data. This is especially true of the stromal remodelling, identified via spatial metrics, which will require robust cross validation to paired patient samples. Technically, the length of time required to generate COJEC-resistant C57 Bl/6 Th-*ALK(F1174L)/MYCN* limits the adaptability of this approach, but syngeneic allografts will be used in the future to provide an alternative approach, partially alleviating this limitation.

## Methods

### *In vivo* experiments

All experiments, including the breeding of transgenic animals, were performed in accordance with the local ethical review panel, the UK Home Office Animals (Scientific procedures) Act 1986, the ARRIVE (Animal Research: Reporting of In Vivo Experiments) guidelines (38) and the UK NCRI guideline (39). All mice used in this study were maintained on a C57 Bl/6J backgraound for >10 generations, and both male and female mice were used in this study. Th-*ALK(F1174L)/MYCN* animals were commenced on COJEC when abdominal palpation of their tumour reached approximately 5mm. After the first cycle, a second cycle was only commenced when palpation revealed tumour progression. This approach was continued until no response to the initial doses of COJEC was seen, at which point, a higher dose of COJEC was started. Low dose COJEC: vincristine 0.125mg/kg, cisplatin 0.5mg/kg, cyclophosphamide 10mg/kg, etoposide 2mg/kg, carboplatin 10mg/kg; high dose COJEC: vincristine 0.125mg/kg, cisplatin 1mg/kg, cyclophosphamide 25mg/kg, etoposide 2mg/kg, carboplatin 15mg/kg. When no tumour response was detected to the higher dose of COJEC, the experiment was terminated and the tumour was deemed resistant. Changes in tumour volume were visualised using MRI on a 7T horizontal bore MicroImaging system (Bruker Instruments) using a 3 cm birdcage coil. Anatomical T_2_-weighted coronal images were acquired through the mouse abdomen, from which tumour volumes were determined using segmentation from regions of interest (ROI) drawn on each tumour-containing slice.

### Statistical analyses

Statistical analysis was performed using the two-tailed Student’s t-test for single comparison, one-way ANOVA or two-way ANOVA for multiple comparisions, with post-hoc Tukey analysis. Survival comparisons were made with Log-rank Mantel Cox test. Differences were considered significant at *p<0.05; **p<0.01; ***p<0.001; ****p<0.0001.

### Multiplex immunofluorescence

Multiplex immunofluorescence staining was performed on FFPE sections using the OPAL protocol (Akoya Bioscences) on the Bond autostainer system (Leica Bond Rx). Sections were subjected to six sequential rounds of staining with each primary antibody followed by a second HRP-conjugated polymer (Leica Novolink Polymer Detection system). Signal amplification was achieved with TSA-Opal fluorophores (Akoya, supplementary table S1). Before the first round and between each round of staining, a heat-induced epitope retrieval step was performed (BOND epitope retrieval solution). After the final round of antibody staining, slides were counterstained with DAPI (Thermo Fisher Scientific) and mounted with ProLong Diamond antifade mounting medium (Thermo Fisher Scientific).

### scRNAseq

Tumour samples were minced and enzymatically digested (Miltenyi Mouse Tumour Dissociation Kit, 130-096-730). Femurs were either crushed or flushed with PBS to release bone marrow. Single cell suspensions were fixed according to Parse Biosciences Evercode^TM^ Cell Fixation v3, allowing samples to be stored and batched after fixation, prior to combinatorial barcoding according to Parse Biosciences Evercode^TM^ WT v3. Sequencing was carried out at a depth of 30,000 reads per cell (Illumina NovaSeq6000).

### Flow Cytometry

Tumours and femours were prepared as for scRNAseq. After filtering, red cell lysis was performed (ACK buffer, Sigma, A1049201) and cells re-suspended in flow cytometry staining buffer (PBS with 2% fetal calf serum and 2mM EDTA) for counting and subsequent immunostaining. Surface antibodies and viability dye (supplementary table S2) were applied for one hour at 4°C, before permeabilization (Foxp3/Transcription Factor Staining Buffer Set, eBioscience 00-5523-00). Cells were then stained with intracellular antibodies for a further hour at 4°C. Cells were fixed (1% paraformaldehyde) and measured on a BD Symphony A5 Cell Analyzer within 72 hours.

### Spatial immunophenotyping Sectioning

Mice tissues were flash frozen and embedded in OCT (ThermoFischerScientific) compound for cryosectioning. 6µm thick sections were obtained using cryostat and mounted into SuperFrost Plus (VWR) charged slides. Sections were stored in -80°C for long term storage until assay was performed.

### Phenocycler-Fusion Staining

Antibodies were purchased from Akoya Biosciences and Leinco Technologies or custom conjugated with Akoya’s custom conjugation kit (7000009), barcodes and protocol as detailed in Supplementary table S3.

Phenocycler-Fusion staining was performed as Akoya’s protocol for Flash Frozen assays as follows. Sections were removed from -80°C and directly laid on top of Drierite (VWR) for 5 minutes to dry. Subsequently sections were immersed in Acetone for 10 minutes for initial fixation. After removal from Acetone sections were let to dry at room temperature for 2 minutes and then immersed in two jars filled with Hydration Buffer (Akoya Biosciences) four times each and incubated 2 minutes each. Following incubation slides were immersed in Pre-Staining Fixing solution (Staining Buffer, Akoya Biosciences,16%PFA) for 10 minutes followed by four immersions in two rounds of Hydration Buffer. Then slides were incubated in Staining Buffer (Akoya Biosciences) for 25 minutes. During incubation time an antibody cocktail was prepared with Staining Buffer and containing S, G, N and J blockers (Akoya Bioscience). Antibodies were added to antibody cocktail as per dilution in Supplementary Table S3, dispensed onto the sections and incubated for 3h at room temperature. Following incubation slides were immersed four times each in two rounds of fresh Staining buffer and then incubated in Post-Staining Fixing Solution (Storage Buffer, Akoya Biosciences, 16% PFA). Slides were then washed three time in PBS and incubated in ice cold Methanol for 5 minutes on ice and followed by three round of PBS washes. A final fixation round was performed with Fixative Solution (Fixative Reagent, Akoya Biosciences, PBS) for 20 minutes at room temperature. A final three round of washes in PBS was performed. Slides were then stored in Storage Buffer at 4°C or immediately processed for Phenocycler run.

### Flow cell assembly and Phenocycler-Fusion run

Slides were incubated in PBS for 10 minutes at room temperature. Flow cell was assembled as per Akoya’s protocol and incubated for 10 minutes in 1X Buffer for Phenocycler plus Additive. A reporter plate was prepared as per Akoya’s instructions and loaded onto the machine. Phenocycler run was launched through Akoya’s Fusion software version 2.1.0.

### Cell classification and spatial analysis

Images generated by Phenocycler-Fusion and OPAL were visualised in QuPath (v0.5.1). Cell segmentation was performed using StarDist extension (v0.5.0). Cell classification was conducted in Qupath using a threshold-based approach, whereby marker positivity was defined using single-measurement classifiers based on mean cellular or nuclear intensity values, as appropriate. Individual marker classifications were subsequently combined using a custom script implementing hierarchical rules to assign composite cell phenotypes based on marker co-expression patterns. Spatial analysis was conducted with MuSpAn v1.2.4 (44).

### Immunohistochemistry

Tumours and bones from Th-*ALK(F1174L)/MYCN* animals were fixed (4% paraformaldehypde) for 48 hours at 4°C. Bones were subsequently decalcified (Osteosoft, Sigma) for one week. After paraffin embedding, all tissue blocks were cut to provide sections of 4µm. Immunohistochemistry was performed for Ly6G (87048, Cell Signaling Technology), Neutrophil Elastase (90120T, Cell Signaling Technology) and Citrullinated Histone H3 (97272T, Cell Signaling Technology), using standard methods, including heat-induced epitope retrieval using citrate buffer (pH6).

### Biomarker assays

Calprotectin degradation mediated via Human Neutrophil Elastase (HNE) was quantified by targeting a specific degradation fragment in a stored plasma using competitive enzyme-linked immunosorbent assay (ELISA) CPa9-HNE, as described previously (1031AE01, Nordic Bioscience) (45). Briefly, CPa9-HNE quantifies a highly specific neoepitope of calprotectin in the S100A9 subunit generated via HNE. Biomarker values outside the quantification range were assigned either the lower or upper limit of quantification, as appropriate. Total calprotectin (S100A8/S100A9) measurements in the plasma were conducted with a commercially available sandwich ELISA (R and D Systems, DS8900).

### Patient samples

Patient-derived material and clinical data were collected under institutional ethical approval, and in accordance with the Declaration of Helsinki. At The Royal Marsden Hospital, patient samples were collected as part of the European ITCC-P4 programme (ITCC-P4 GmbH Paediatric Preclinical Proof of Concept Platform; UK IRAS ID 243123; local CCR study ID 4892) with infomed consent from all patients or their legal guardians.

## Declarations

### Availability of data and materials

The datasets supporting the conclusions of this article will be made available in the xxx repository (http://)

### Funding

This study was funded by the Cancer Cluster programme within the Medical Research Council National Mouse Genetics Network (MC_PC_21042). ERT is supported by a US Department of Defence Award (HT9425-24-1-0439, CA230885). AL is supported by The Team Luke Foundation (RFR246X). KB is supported by the Medical Research Council National Mouse Genetics Network (MC_PC_21042). SM and VT are supported by The Children and Young People’s Cancer Association, Little Princess Trust, UK (LPT2024A30). GM was supported by a Neuroblastoma UK award (NBUKCheMartin25). LC, BMC and KG were supported by a Cancer Research UK Programme Grant (A78278). LC is funded by ICR/HEFCE.

### Authors’ contributions

ERT conceptualized the study. Experiments were performed by ERT, AL, KB, BMC, KG, FL, SM, VT, NSH, GE and GM. Bioinformatic analysis and interpretation was carried out by ERT, AL, JH, DR and MB. Patient samples were provided by ET, JP, PA, SG. CPa9-HNE assay was carried out by NW and MP. Project supervision was undertaken by MY, NB, TG, SG and LC. Funding and resources were acquired by LC. ERT wrote the manuscript.

## Acknowledgements

ERT would like to acknowledge Mr Tony Rogers for clinical sample coordination, the Flow Cytometry Facility, Genomics Facility and The Biological Services Unit at The Institute of Cancer Research for their support of this study, and the patients, their families and staff from The Royal Marsden Hospital who generously participated in this research.

## Notes

### Competing Interest Statement

The authors have declared no competing interest.

