## Supplementary Figures for "A COJEC-chemotherapy resistant model of Th-*ALK(F1174L)/MYCN* neuroblastoma offers insights into tumour immune evasion and development of the bone marrow metastatic niche"

Supplementary figure one

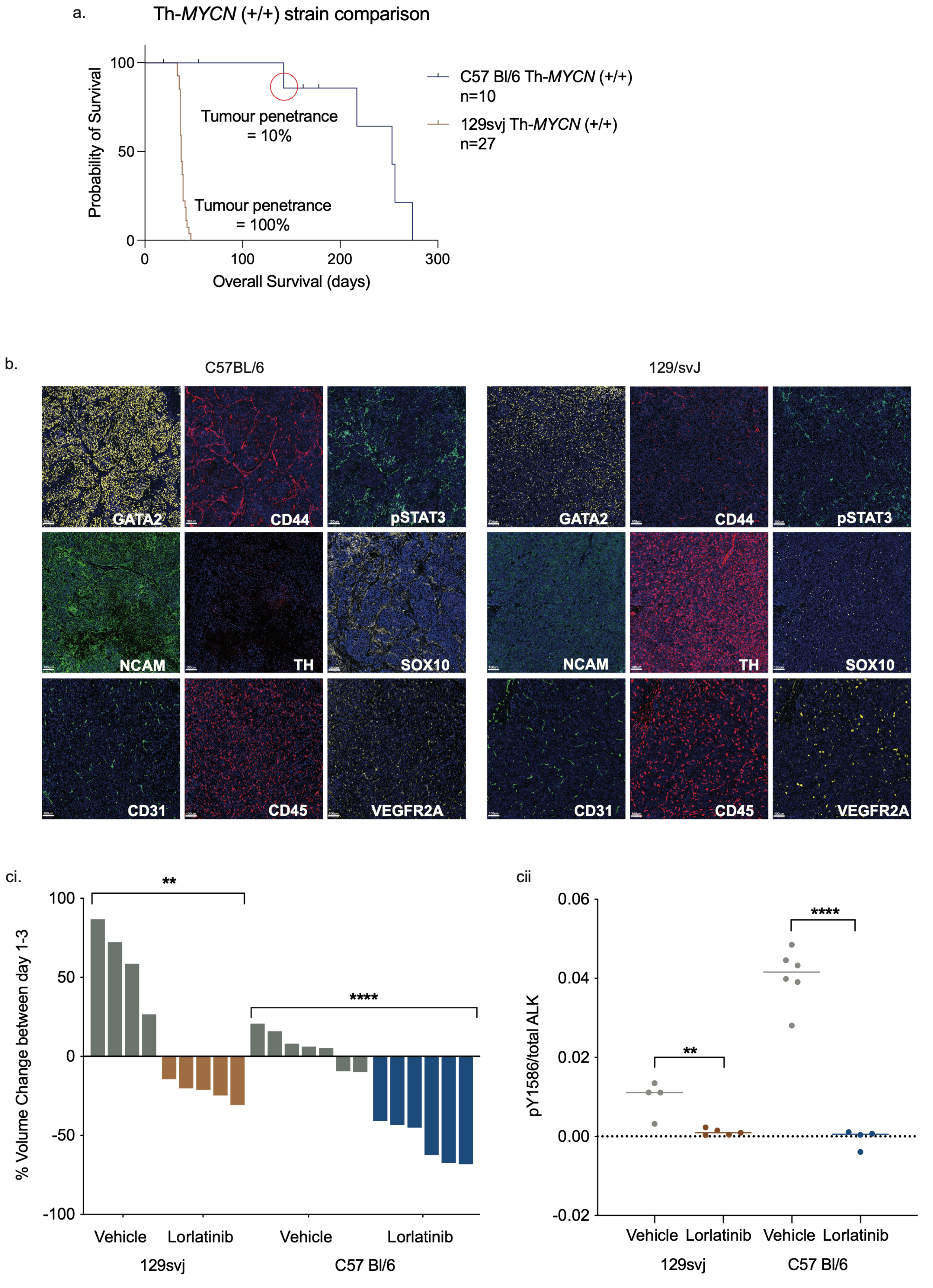

Supplementary figure two

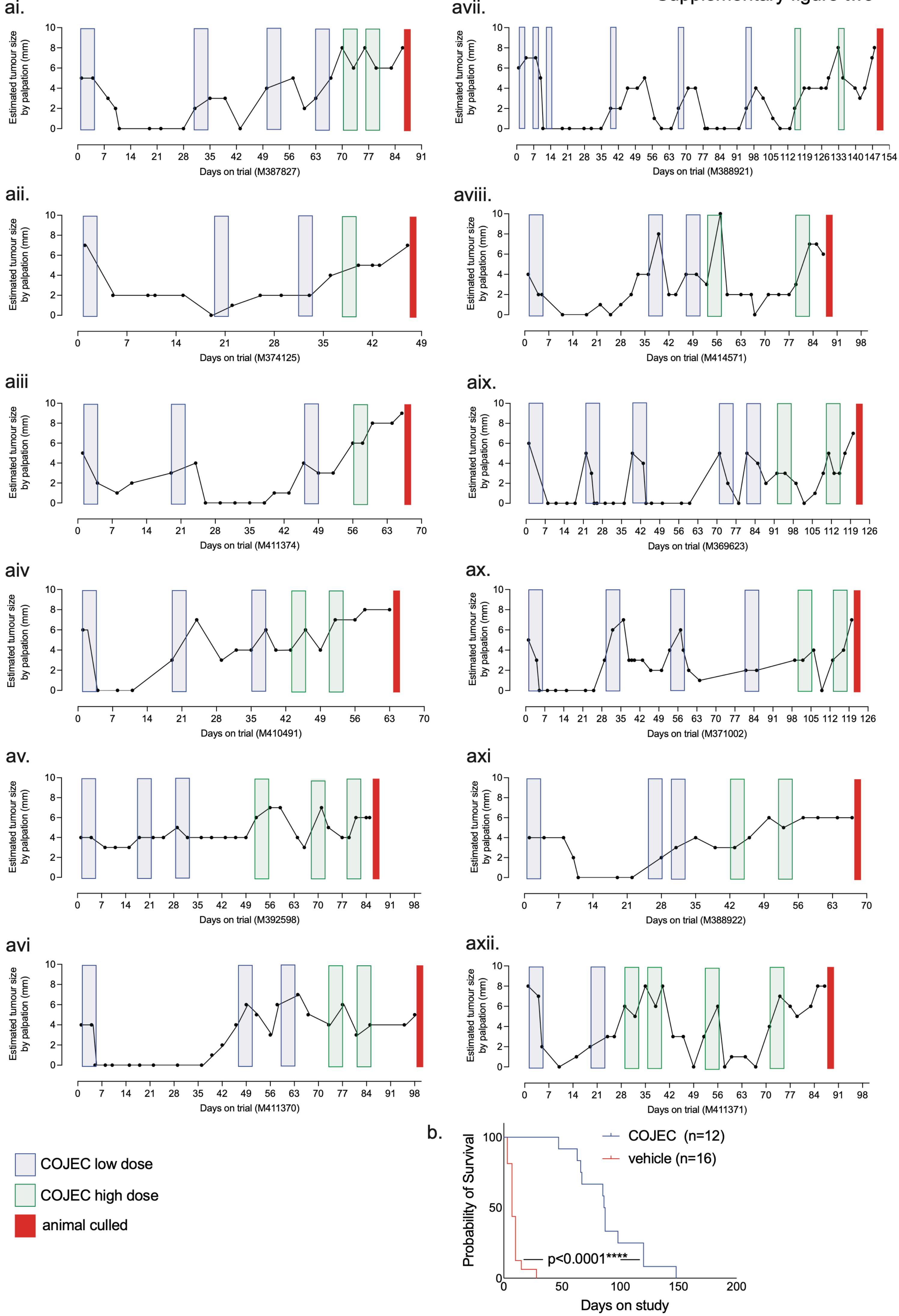

### Supplementary figure three

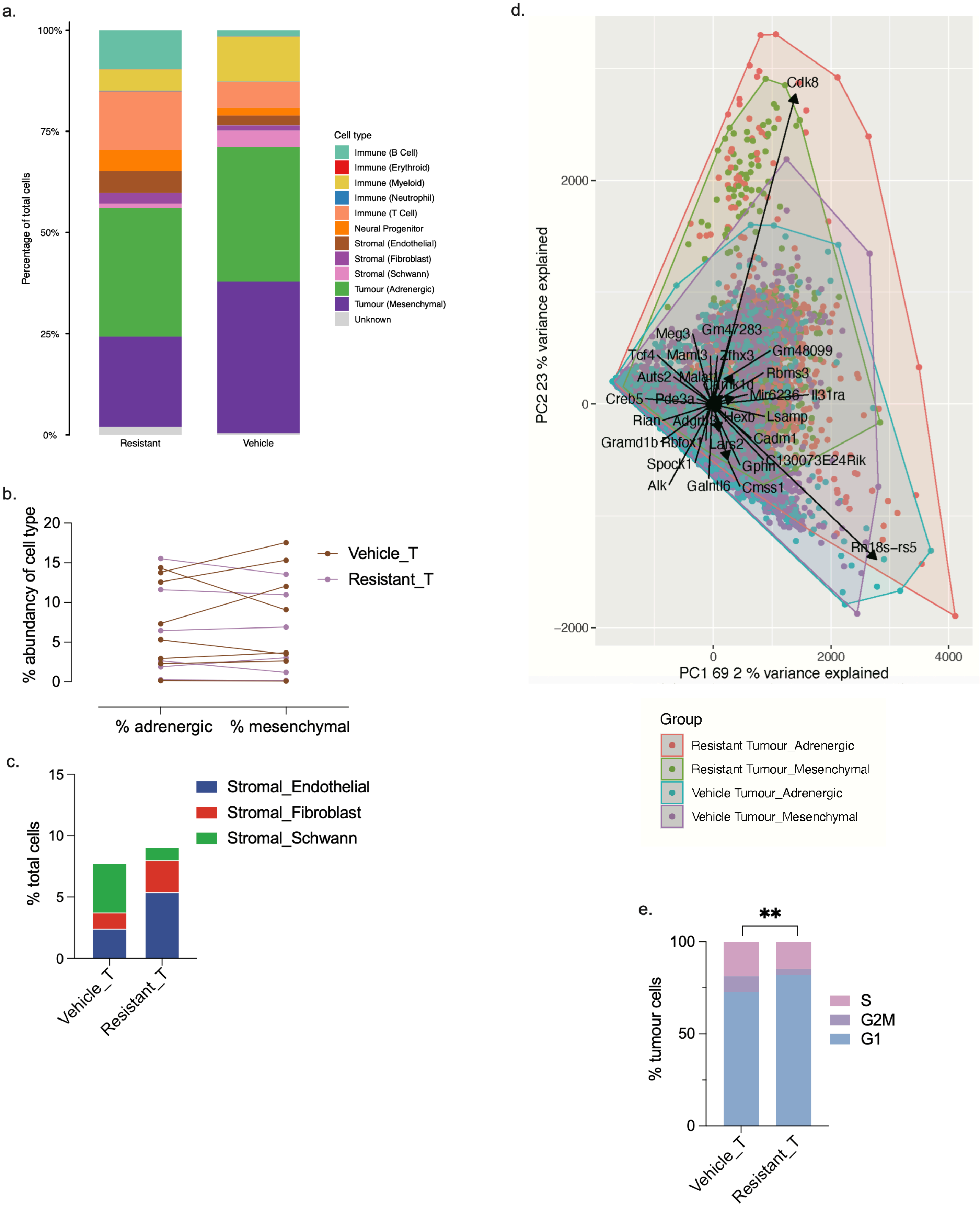

Supplementary figure four

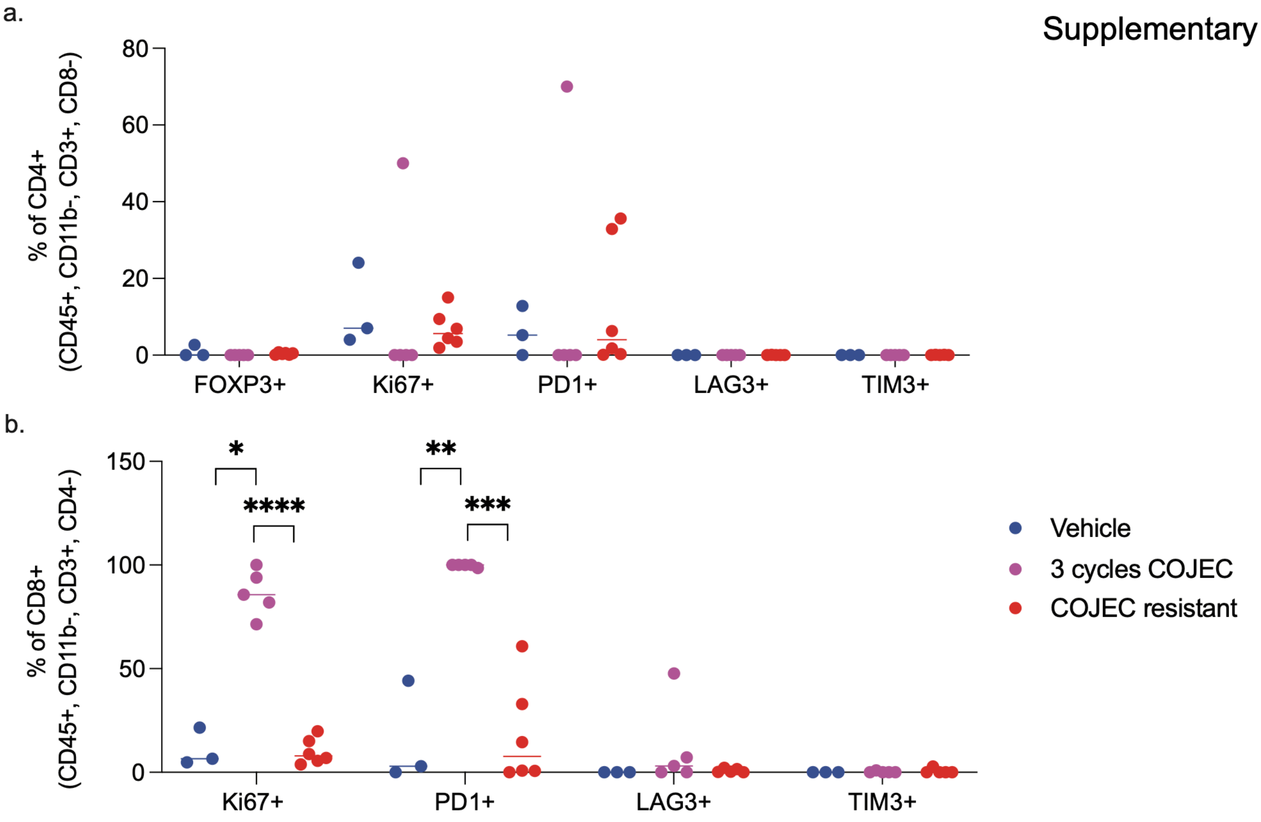

### Supplementary figure five

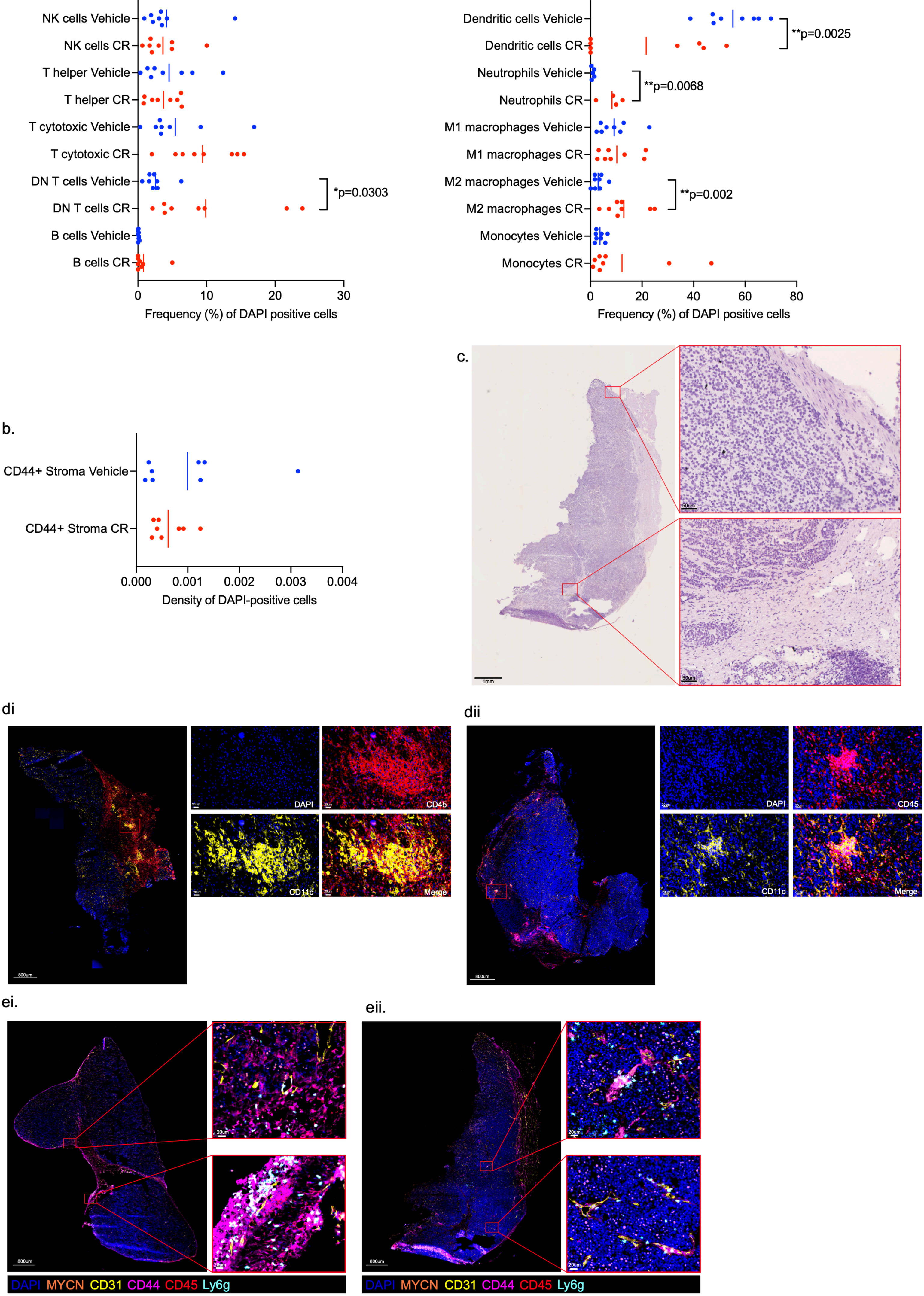

#### Supplementary figure six

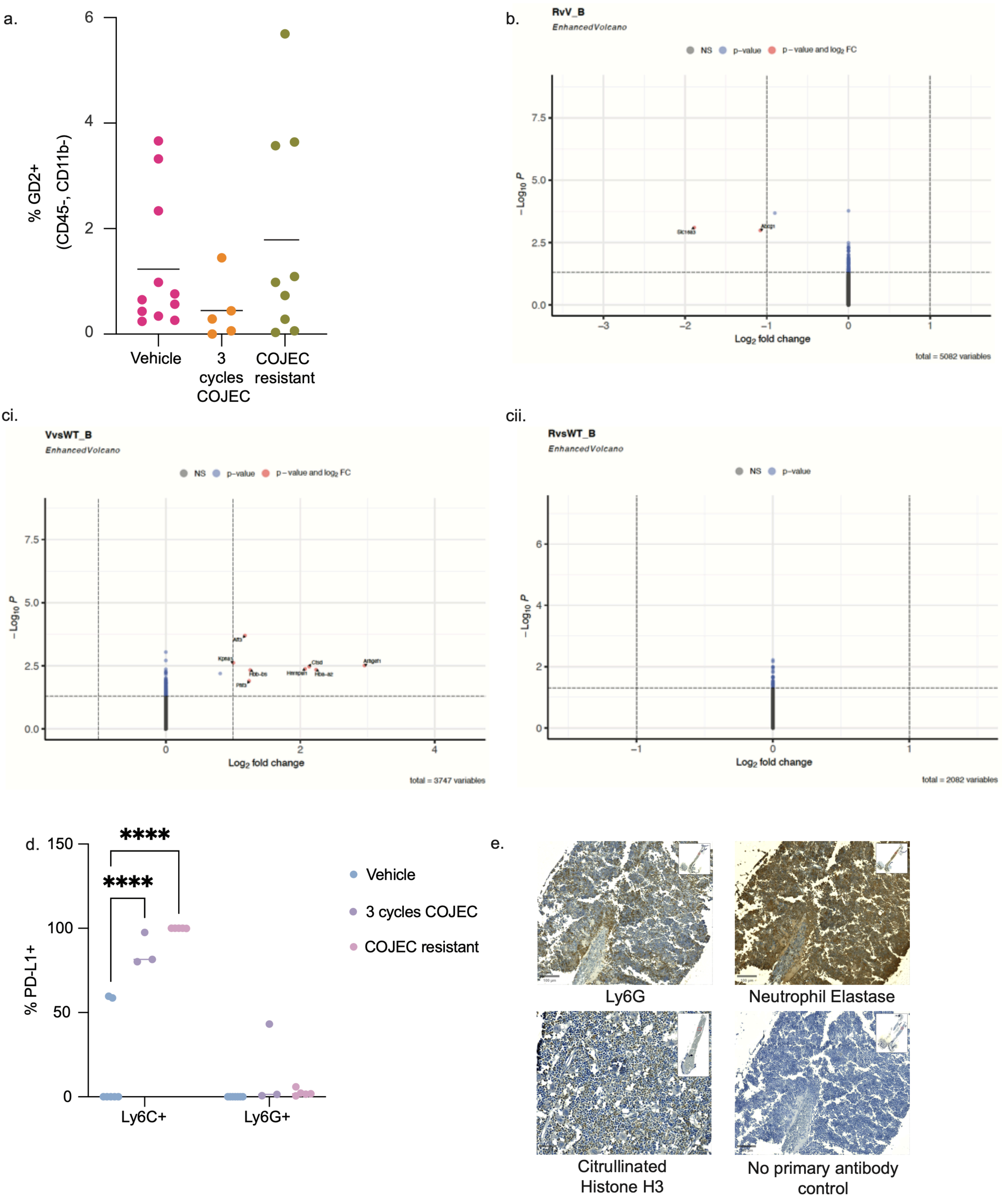

#### Supplementary figure seven

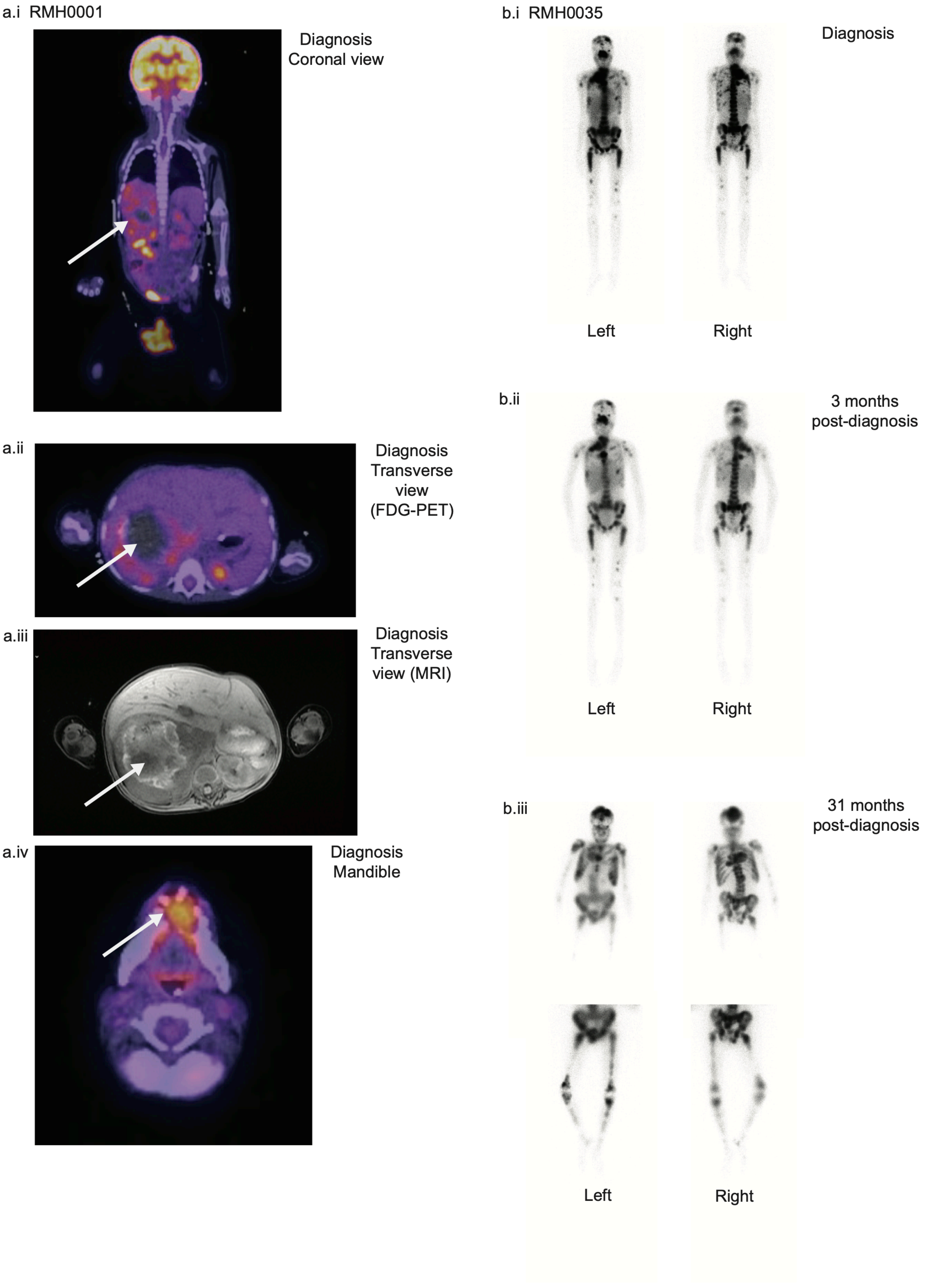
