## Supplementary Figure Legends for "A COJEC-chemotherapy resistant model of Th-*ALK(F1174L)/MYCN* neuroblastoma offers insights into tumour immune evasion and development of the bone marrow metastatic niche"

Supplementary Figure One

a. Kaplan Meier survival plot of C57 Bl/6 versus 129 svj Th-*MYCN* (+/+) mice. One C57 Bl/6 mouse developed an abdominal neuroblastoma (circled on plot). b. Immunofluoresecence (Opal) example staining of ethical endpoint tumours from C57 Bl/6 and 129svj Th-*ALK(F1174L)/MYCN*. ci. Waterfall plot depicting percentage tumour volume change between day 1 and day 3 in Th-*ALK(F1174L)/MYCN* mice treated with either Lorlatinib (10mg/kg) or vehicle control. cii. ALK and pALKY1586 immunoassay of tumours samples taken from mice in ci. Assay run according to published protocol (1).

Supplementary Figure Two

ai-xii. Tumour palpation data for all animals included in the COJEC-resistance study. The blue rectangular box indicates administration of a low-dose cycle of COJEC, the green box indicates administration of a high-dose cycle of COJEC, the red box indicates the end of the study (ethical end point). b. Kaplan-Meier survival plot of animals treated with cycles of COJEC chemotherapy (individually depicted in a), versus animals treated with vehicle control.

Supplementary Figure Three

a. scRNAseq cell type percentage of total cells in COJEC-resistant and vehicle tumour samples. b. percentage adrenergic and mesenchymal tumour cells in individual tumours. c. Endothelial / Fibroblast / Schwann Stroma percentage total cells in vehicle and COJEC-resistant tumour samples. d. Principal Component Analysis Bi-plot identifying Cdk8 within Resistant Tumour Adrenergic group. e. cell cycle analysis of tumour cells in vehicle and COJEC-resistant tumours.

Supplementary Figure Four

Flow cytometry analysis of tumour samples, showing a. CD4+ T-cell subsets and b. CD8+ T-cell subsets.

Supplementary Figure Five

Frequency of a.i. lymphoid cells and, a.ii. myeloid cells in Vehicle and COJEC-resistant (CR) Th-*ALK(F1174L)/MYCN* C57 Bl/6 tumours according to analysis of phenocycler sections. b. Density of stroma (CD44) in Vehicle and CR tumours. c. Haematoxylin and eosin staining (post-phenocycler antibody staining) of section depicted in figure 4c. d.i. and d.ii. Examples of COJEC-resistant tumours with dendritic cell (CD11c) clusters. e. Example of i. Vehicle tumour section Neighbourhood 1, demonstrating a “vascular (CD31, endothelial cells) immune enriched” region, and ii. a COJEC-resistant tumour Neighbourhood 4, demonstrating a “vascular-stroma immune depleted” region, with the exception of neutrophils (Ly6g).

Supplementary Figure Six

Flow cytometry analysis of bone marrow from crushed/flushed femurs to measure GD2+ (CD45-, CD11b-) cells. b. scRNAseq differential gene expression (DEG) analysis of bone marrow samples between vehicle and resistant samples. c. scRNAseq paired DEG analysis from bone and tumour samples from i. vehicle-treated mice and ii. COJEC-resistant mice. d. Flow cytometry analysis of PDL1 expression on Ly6C and Ly6G MDSC cells from bone marrow samples. e. Example immunohistochemistry of FFPE femurs from COJEC-resistant mouse for markers of neutrophils (PMN-MDSC) (Ly6G) and neutrophil NETosis (Neutrophil Elastase and Citrullinated Histone H3).

Supplementary Figure Seven

a. Staging fluorodeoxyglucose Positron Emission Tomography (FDG PET) scan for patient RMH0001, demonstrating a.i. (coronal view), a.ii. (sagittal view), a.iii (corresponding MRI) large right inhomogeneous suprarenal neuroblastoma, partly calcified, with local effect on liver and kidney and left para-aortic nodal disease. a.iv. 16mm FDG-avid soft tissue mass arising from the middle mandible, with focal expansion of the medullary space, splaying of the midline dentition and thinning of the overlying superficial and deep bone cortex (SUV max 3.4). In addition, patient reported to have multiple permeative osseous deposits associated with mild FDG uptake including proximal left humeral diaphysis with possible pathological fracture. Involvement of the knees, ankles and lateral 9^th^ rib (imaging not available). b. Meta-iodobenzylguanidine (mIBG) scans for patient RMH0035 demonstrating chemotherapy-refractory disseminated skeletal disease. b.i. Diagnostic mIBG demonstrating mIBG-avid skeletal deposits, associated with mixed lytic / sclerotic changes on recent CT (images not shown). b.ii. mIBG prior to PARC Trial enrolment showing no significant change or avidity of disseminated skeletal disease. b.iii. Reassessment mIBG showed enlargement of masses associated with left rib, with associated increase in tracer avidity. Disseminated skeletal disease with multiple pathological fractures.
